# Layer-dependent activity in the human superior colliculus during working memory

**DOI:** 10.1101/2022.12.06.518975

**Authors:** Danlei Chen, Philip A. Kragel, Tor D. Wager, Lawrence L. Wald, Marta Bianciardi, Ajay B. Satpute, Karen S. Quigley, Lisa Feldman Barrett, Yuta Katsumi, Jordan E. Theriault

## Abstract

We examined the superior colliculus (SC) with ultra-high resolution 7-Tesla fMRI during an N-back working memory task. We observed both increased BOLD signal intensity and functional connectivity that followed a layer-dependent pattern predicted from anatomical connections between SC and other brain structures important for visual processing, motor control, and executive function. Our results highlight a role for the human SC in cognitive functions that usually associated with the cerebral cortex.

---

The superior colliculus (SC) is a laminated structure in the vertebrate midbrain with three distinct anatomical layers along a dorsolateral to ventromedial axis (May, 2006). Anatomical and behavioral evidence from non-human vertebrates indicates that the superficial layers are primarily visual sensory in nature, whereas the intermediate and deep layers are multisensory and contribute to both skeletomotor and visceromotor control (for review, see Gandhi & Katnani, 2011; Liu et al., 2022), coordinating fast motor outputs in threatening and appetitive encounters (see Isa et al., 2021). Research in non-human vertebrates further supports a role for the intermediate and deep layers of SC in selective attention (e.g., Lovejoy & Krauzlis, 2010) and perceptual decision-making (e.g., Jun et al., 2021; for review, see Basso et al., 2021). Research in humans, however, has primarily examined the SC in visual stimulation and visually guided behaviors (e.g., Krebs et al., 2010; Loureiro et al., 2017; Wang et al., 2020). The present study investigated the human SC in a working memory task that required visual perception, motor response, and executive function, thereby bridging the gap between human and non-human vertebrate research on the SC.

Participants (N = 88) completed 12 blocks of an N-back working memory task in which they were presented with a sequence of letters and were asked to press a button when the current letter matched the letter in the prior trial (1-back, low cognitive load condition) or two trials back (3-back, high cognitive load condition). Successful performance involved the processing of visual sensory signals, maintenance of the information across trials (i.e., working memory), control of motor systems to support button presses, and goal-oriented executive control and coordination of these functions. As a motivated performance task, the N-back task engages changes in autonomic control of visceromotor activity (e.g., Mandrick et al., 2016), and these changes were confirmed by the behavioral and peripheral physiological results in the present task (see Fig. S1 for behavioral and physiological changes). Based on non-human vertebrate research, we hypothesized that the SC would show an increase in blood-oxygen level-dependent (BOLD) signal intensity for both cognitive load conditions compared to baseline (i.e., fixation). Further, we hypothesized that SC would show a greater increase in BOLD intensity for higher cognitive load 3-back condition relative to the 1-back condition (while visual stimulation was identical, and the number of motor responses did not significantly differ across conditions; see Fig. S1). We also hypothesized the SC would show a greater increase in BOLD intensity in the intermediate and deep layers (ventromedial SC) compared to the superficial layers (dorsolateral SC), because the intermediate and deep layers of SC have been implicated in the integration of multiple functions that are required for performing the N-back task, whereas the superficial layers have been implicated only in visual perception. Finally, we investigated whether the superficial layers would show strong connectivity with other brain structures important for visual processing during the entire task, whereas the intermediate and deep layers of SC would show strong connectivity with structures important for multiple functions related to the present task beyond vision, including visceromotor control and executive function.

We spatially aligned voxels corresponding to the anatomical location of the superior colliculus into a common space across participants. As hypothesized, the entire SC showed a significant increase in BOLD intensity in 3-back > baseline (*t*_*(76)*_ = 11.33, *P* < .001), 1-back > baseline (*t*_*(82)*_ = 7.20, *P* < .001), and 3-back > 1-back condition (*t*_*(71)*_ = 5.75, *P* < .001) (see the voxel-wise comparison in Fig. 1A). A visual inspection of the SC contrasts confirmed that the BOLD intensity was greater in the ventromedial subregion (the deep layers) compared to the dorsolateral subregion (the superficial layers). To quantify this difference, we estimated the layer structure of the superior colliculus (i.e., voxel depth; Fig. 1B) and correlated voxel depth with voxel-wise BOLD intensity (Fig. 1C). We then estimated a linear mixed effects model that used cognitive load, voxel depth, laterality, and their full interactions as fixed effects. As hypothesized, the SC showed a significant increase in BOLD intensity in the 3-back compared to 1-back conditions (*F*_*(1, 81.78)*_ = 14.84, *P* < .001) and in the intermediate and deep layers compared to superficial layers (*F*_*(1, 77.48)*_ = 24.62, *P* < .001) (Fig. 1D; see Online Methods for anatomical, preprocessing, and modeling details).

**Fig. 1.**
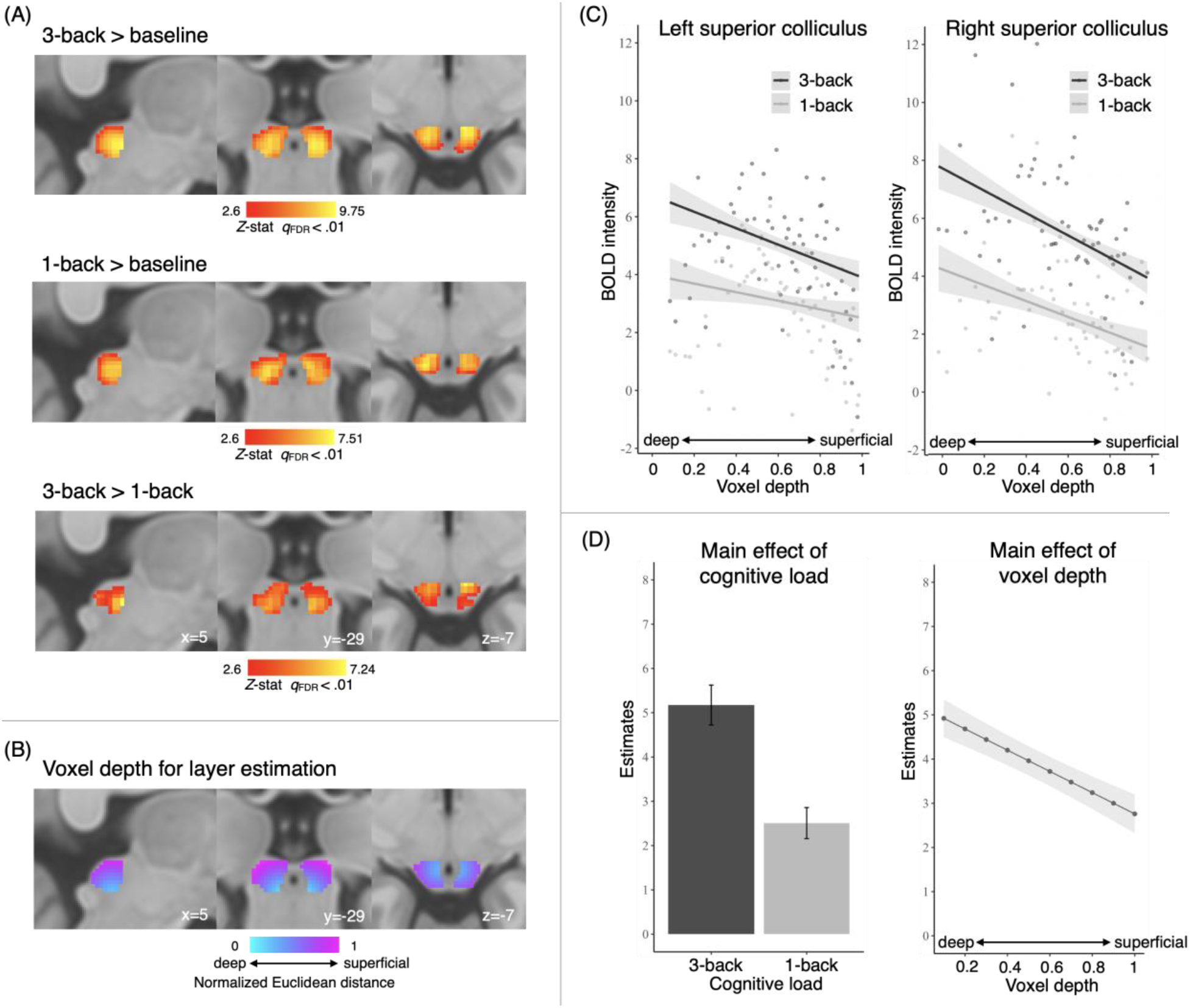
Differences in BOLD intensity of the SC during an N-back working memory task. (A) Group-level univariate contrasts of the SC in 3-back > baseline, 1-back > baseline, and 3-back > 1-back (see Fig. S2 for whole-brain contrasts). Importantly, visual stimulation was identical across the cognitive load conditions, and the number of motor responses did not significantly differ across conditions (see Supplementary Material for behavioral results); therefore, any difference in BOLD intensity in the SC between the cognitive load conditions was not likely to be a product solely from any differences in visuomotor processing. Colored voxels indicate one-tailed group-level comparison in *Z* statistics using first level (participant-level) means, thresholded at a voxel-wise cutoff of *q*_FDR_ < .01 within the group-specific SC mask (see Online Methods and Fig. 1B). The SC univariate contrasts are available [online](https://osf.io/mhtce/?view_only=0cd2e83463be4df8bda9537db5173aa3). (B) Voxel depth estimates of the SC layer structure. Euclidean distance was calculated from each voxel within the SC mask to an anchor location ventromedial to the SC on each lateral side (anchor location: left side: x=-3, y=-30, z=-10; right side: x=3, y=-30, z=-10). A voxel depth closer to 0 represents the deeper SC layers and a voxel depth closer to 1 represents the superficial layers. (C) Voxel-wise relationships between group-averaged SCBOLD intensity (y-axis) and cognitive load (3-back/1-back), voxel depth (0-1), and laterality (left/right). Linear regressions of BOLD intensity were fit across voxel depth in each cognitive load condition. (D) Estimates of marginal means for significant main effects of cognitive load and voxel depth. The effects of cognitive load, voxel depth (rank-ordered and discretized into 10 bins), laterality, and full interactions were submitted to a linear mixed effects model (see Online Methods), which identified the main effects of cognitive load (3-back > 1-back; *F*_*(1, 81.78)*_ = 14.84, *P* < .001) and voxel depth (deep > superficial; *F*_*(1, 77.48)*_ = 24.62, *P* < .001). All other main effects and interactions were non-significant (*P* > .05). See full model output in Table S1. Error bars indicate ± 1 standard error across participants.

We investigated the functional connectivity between the SC and the cerebral cortex by estimating functional connectivity gradients (see e.g., Hong et al., 2020; Katsumi et al., 2021), computed as the temporal correlation of BOLD intensity fluctuations between 377 bilateral SC voxels and 1000 cortical parcels (Schaefer et al., 2018), using the residual timeseries after regressing out stimulus-evoked changes in BOLD intensity (Fig. 2A). The dominant gradient (i.e., Gradient 1; accounted for 14.69% of variance) revealed a connectivity pattern that transitioned along the dorsolateral-to-ventromedial axis of SC (Fig. 2B) and was strongly correlated with voxel depth (Spearman’s rank correlations: *ρ* = .79, *P* < .001), suggesting that Gradient 1 corresponded with the anatomical layers within the SC. Additional gradients were either not readily interpretable according to anatomical referents (e.g., Gradient 2, see Fig. S3) or did not account for an appreciable portion of variance in SC-cortical functional connectivity.

**Fig. 2.**
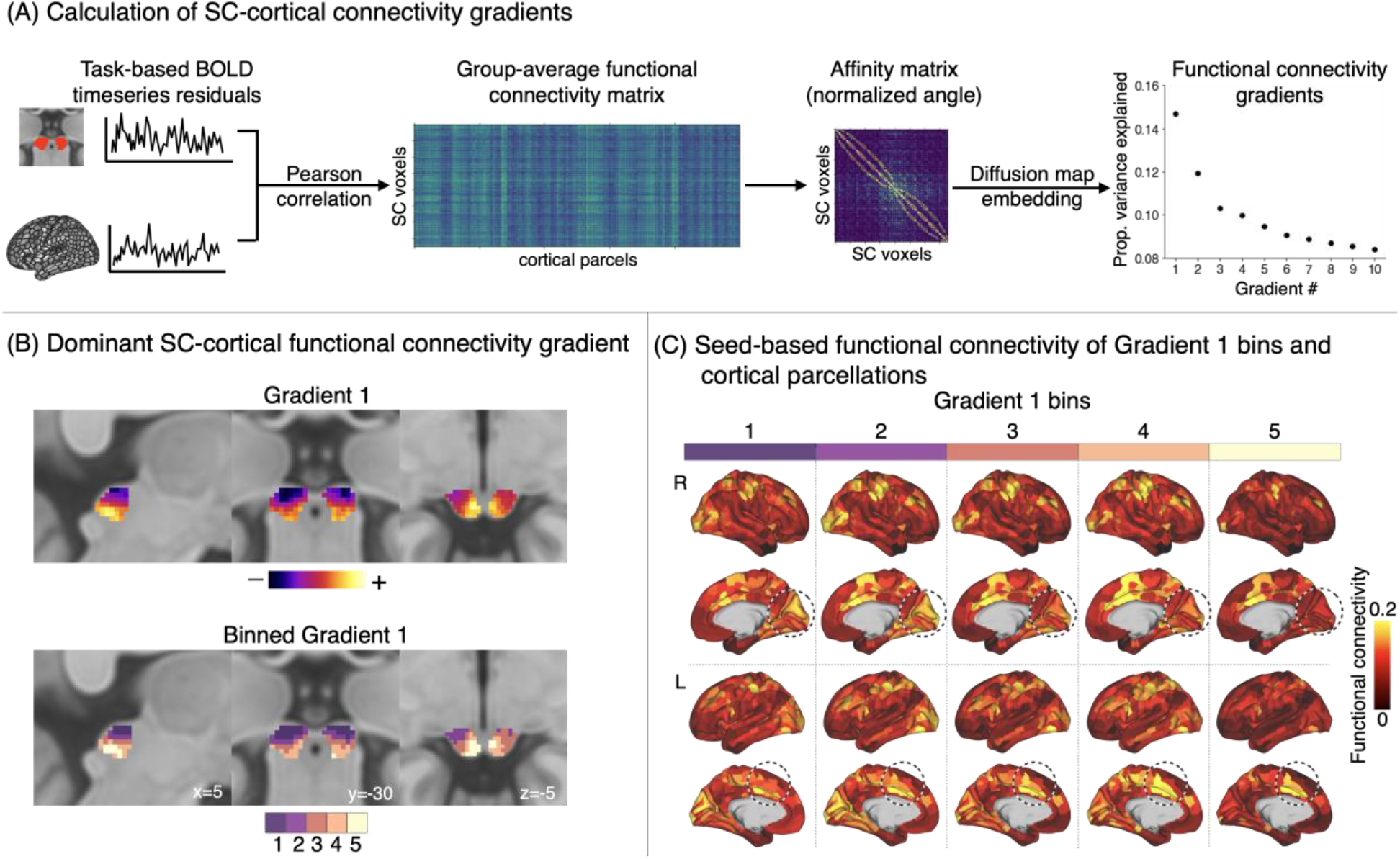
Functional connectivity organization of the SC during an N-back working memory task. (A) Schematic diagram of the functional connectivity gradient analysis. Inputs were participant-level residual BOLD timeseries from the univariate analysis of the N-back working memory task data. In each participant, we calculated a functional connectivity matrix based on Fisher Z-transformed Pearson’s correlation coefficients between each of the 377 voxels within the SC and 1000 spatially discontiguous parcels of the cerebral cortex (Schaefer et al., 2018). To characterize the organization of functional connectivity with the cortical parcels in the superior colliculus, we converted the group-averaged functional connectivity matrix to a non-negative square symmetric affinity matrix expressed as normalized angle similarity. Each cell in this affinity matrix (377 × 377) represented the magnitude of similarity in the pattern of functional connectivity with 1000 cortical parcels between a given pair of SC voxels. The affinity matrix was then used as input for diffusion map embedding, a non-linear approach for dimension reduction, which yielded a set of connectivity gradients representing the dominant dimensions of spatial variation across all voxels in the superior colliculus, in terms of their functional connectivity with the cortical parcels (see Online Methods). (B) The dominant SC functional connectivity gradient (Gradient 1). Gradient 1 showed a gradual transition in functional connectivity pattern along the dorsolateral-ventromedial axis. The proximity of colors indicates a greater similarity of connectivity patterns with the cerebral cortex. Gradient 1 was divided into five subregions based on the percentile ranks of voxel-wise gradient values, where bin 1 was located at dorsolateral SC (corresponding to superficial layers) and bin 5 was located at ventromedial SC (corresponding to deep layers). The SC gradient maps are available [online](https://osf.io/mhtce/?view_only=0cd2e83463be4df8bda9537db5173aa3). (C) Seed-based functional connectivity between SC gradient bins and cortical parcels. Each gradient bin was used as a seed in a functional connectivity analysis with all 1000 cortical parcels. Dashed circles around the anterior mid cingulate cortex and visual cortex highlight cortical areas that showed strong cross-bin variations in functional connectivity by visual inspection. For ease of comparison across cortical maps, the range of color bar was held fixed across cortical maps, and the exact connectivity values are available [online](https://osf.io/mhtce/?view_only=0cd2e83463be4df8bda9537db5173aa3). Abbreviation: SC: superior colliculus.

We explored the pattern of connectivity in Gradient 1 in more detail by parsing it into five bins based on the percentile ranks of voxel-wise gradient values (representing SC voxel depth and layers; Fig. 2B). We estimated the seed-based functional connectivity for each Gradient 1 bin to all cortical parcels (Fig. 2C). As hypothesized, cortical parcels involved in vision (e.g., primary visual cortex) showed the strongest linear decrease in functional connectivity with SC from the superficial to deep layers, whereas parcels involved in memory, visceromotor and skeletomotor control, and executive function (e.g., parahippocampal cortex, anterior mid cingulate cortex, ventral anterior insula) showed the strongest linear increase in functional connectivity with SC from the superficial to deep layers (Fig. 3A).

**Fig. 3.**
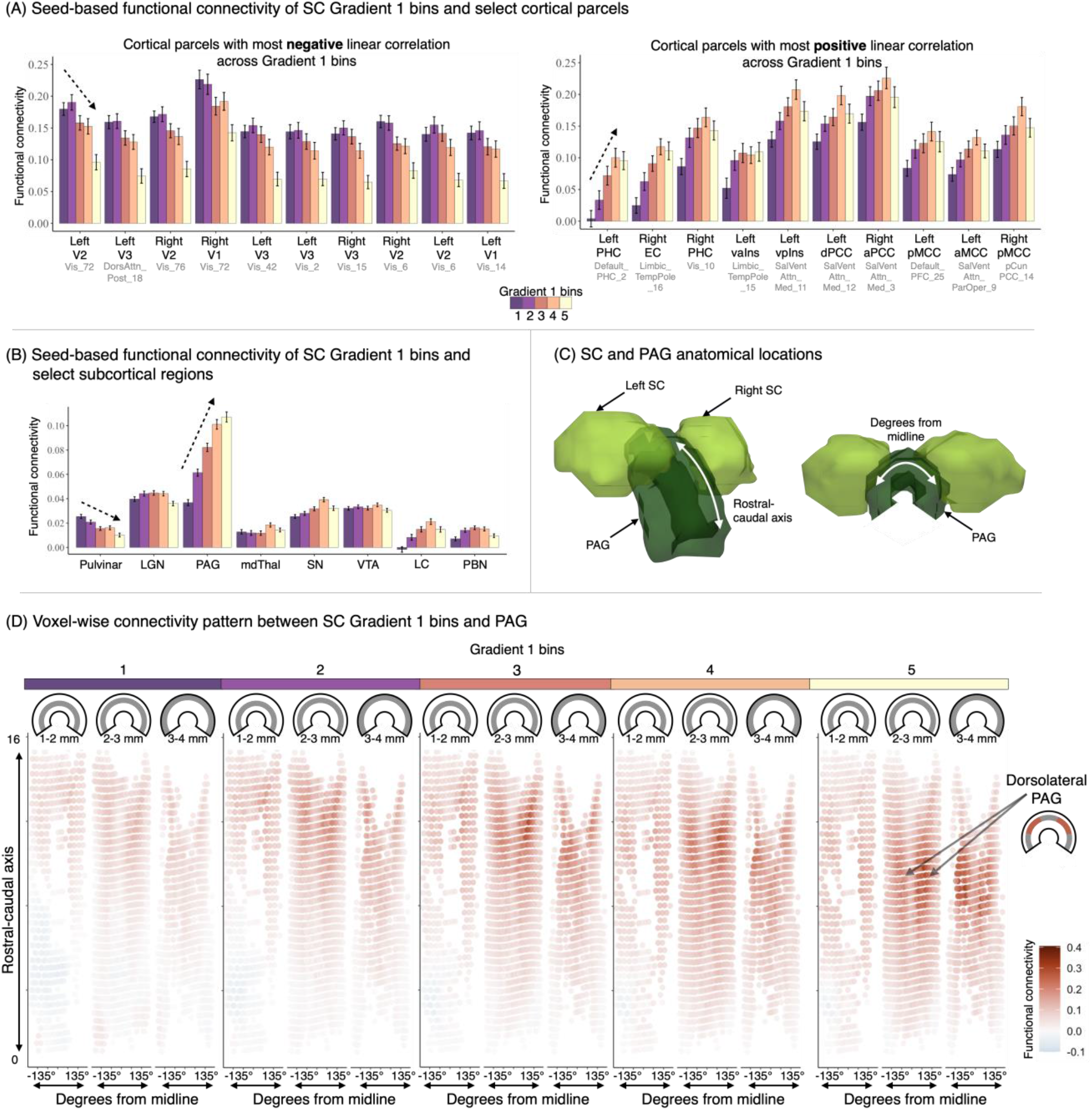
(A) Seed-based functional connectivity between SC Gradient 1 bins and select cortical parcels. Cortical parcels were selected based on the rank order of parcels with the most negative and the most positive linear trends in functional connectivity across Gradient 1 bins (1^st^ to 10^th^ in order from left to right on the x-axis). Parcels with strongest linear decrease (i.e., strongest functional connectivity with bins 1 and 2; corresponding to the dorsolateral superior colliculus, near superficial layers) include multiple regions in the visual cortex, while parcels with strongest linear increase (i.e., strongest connectivity in bins 4 and 5; corresponding to the ventromedial superior colliculus, near deep layers) include regions important for complex cognitive functions as well as visceromotor control. Error bars indicate ± 1 standard error across participants. X-axis labels in black denote the anatomical label of each parcel; the original functional label of each parcel is shown in grey (Schaefer et al., 2018). (B) Seed-based functional connectivity between SC Gradient 1 bins and select subcortical regions. While the connectivity gradients were derived with respect to the cortex, we additionally investigated their relations with other subcortical regions whose putative functions and anatomical connections with the SC were relevant to the present task (see Online Methods for details). In Gradient 1, consistent with the results in cortical parcels, a linear decrease was observed in a subcortical region associated with visual processing (pulvinar), and a linear increase as observed in a region important for visceromotor regions and executive functions (e.g., PAG). Error bars indicate ± 1 standard error across participants. (C) A three-dimensional depiction of the SC and PAG location. The SC (light green) lies immediately dorsal to the PAG (dark green), a horseshoe-shaped structure. The PAG shares rich connections and functional similarities with the SC (Carrive & Morgan, 2012) and was found to be functionally involved in the present task (see studies using overlapping data as the present study with a focus on PAG; Fisherbach et al., submitted; Kragel et al., 2019). (D) Seed-based functional connectivity between SC Gradient 1 bins and PAG voxels. Each panel displays three radial slices of the PAG cylinder (e.g., 1-2 mm). The y-axis illustrates the rostro-caudal position of the PAG (mm). The x-axis represents +/- 135° from the brain midline. Across the Gradient 1 bins, bin 5 (ventromedial superior colliculus, near deep layers) showed strong functional connectivity to the dorsolateral column of PAG, consistent with previous anatomical connectivity evidence (Carrive & Morgan, 2012). See Fig. S4 for voxel-wise connectivity between gradient bins and several other subcortical regions (pulvinar, SN, VTA, and PAG). Abbreviation: SC: superior colliculus; V1: primary visual cortex; V2: secondary visual cortex; V3: tertiary visual cortex; PHC: parahippocampal cortex; EC: entorhinal cortex; vaIns: ventral anterior insula; vpIns: ventral posterior insula; dPCC: dorsal posterior cingulate cortex; aPCC: anterior posterior cingulate cortex; pMCC: posterior mid cingulate cortex; aMCC: anterior mid cingulate cortex; LGN: lateral geniculate nucleus; PAG: periaqueductal grey; mdThal: mediodorsal thalamus; SN: substantia nigra; VTA: ventral tegmental area; LC: locus coeruleus; PBN: parabrachial nucleus.

A similar pattern of seed-based functional connectivity was observed between each Gradient 1 bin and select subcortical ROIs chosen based on their monosynaptic anatomical connections with the SC and their functional relevance to the working memory task (see Online Methods for selection criterion). As hypothesized, a subcortical structure involved in vision (i.e., pulvinar) showed the strongest linear decrease in functional connectivity with SC from superficial to deep layers, whereas brainstem structures as part of the dopaminergic and norepinephrine pathway (substantia nigra and locus coeruleus) and a midbrain structure involved in visceromotor control (periaqueductal grey (PAG)) showed the strongest linear increase in functional connectivity with SC from the superficial to deep layers (Fig. 3B).

Furthermore, we estimated the voxel-wise connectivity pattern between the SC and PAG given their close physical proximity, rich anatomical connections, and functional similarities (Carrive & Morgan, 2012). We found the strongest connectivity between SC deep layers and the dorsolateral column of the PAG, a PAG subregion that has been implicated in freezing and flight behaviors as well as the accompanying sympathetic responses (Bittencourt et al., 2004) (Fig. 3D).

The layer-dependent pattern of the SC BOLD intensity we observed in SC BOLD intensity was replicated using a data-driven approach that showed distinct connectivity patterns of the superficial and deep layers of the SC, such that the superficial layers of SC showed strong functional connectivity with cortical and subcortical structures important for visual processing, while deep layers of SC showed strong functional connectivity with cortical and subcortical structures important for a more diverse set of functions, including visceromotor control and executive function. These results are consistent with the well-established anatomical and functional evidence in non-human vertebrates that the superficial layers are exclusively visual, and in contrast, multimodal sensory, motor, and cognitive functions are localized in the intermediate and deep layers (Liu et al., 2022).

Taken together, these findings, for the first time in humans that we know of, suggest that the SC is important for more than just visual processing and visuomotor behaviors, consistent with an emerging perspective in non-human primates that the SC is part of the neural circuity supporting cognitive functions (see Basso et al., 2021). Furthermore, these findings indicate that a working memory task, which is a standard measure of cognitive control, involves more of the brain than just the cerebral cortex. Importantly, our result, for the first time in humans, identified a task-based functional connectivity pattern between the deep layers of the SC and the dorsolateral column of PAG, consistent with the fine-grained and well-replicated tract-tracing evidence in non-human vertebrates (Carrive & Morgan, 2012). This result indicates that the regulation of the body (e.g., visceromotor control) is important not just in life-threatening situations but also in cognitive tasks and presents a future opportunity to investigate a possible role of SC in visceromotor control through its connection with the PAG.

## Supplementary Material

**Fig. S1.**
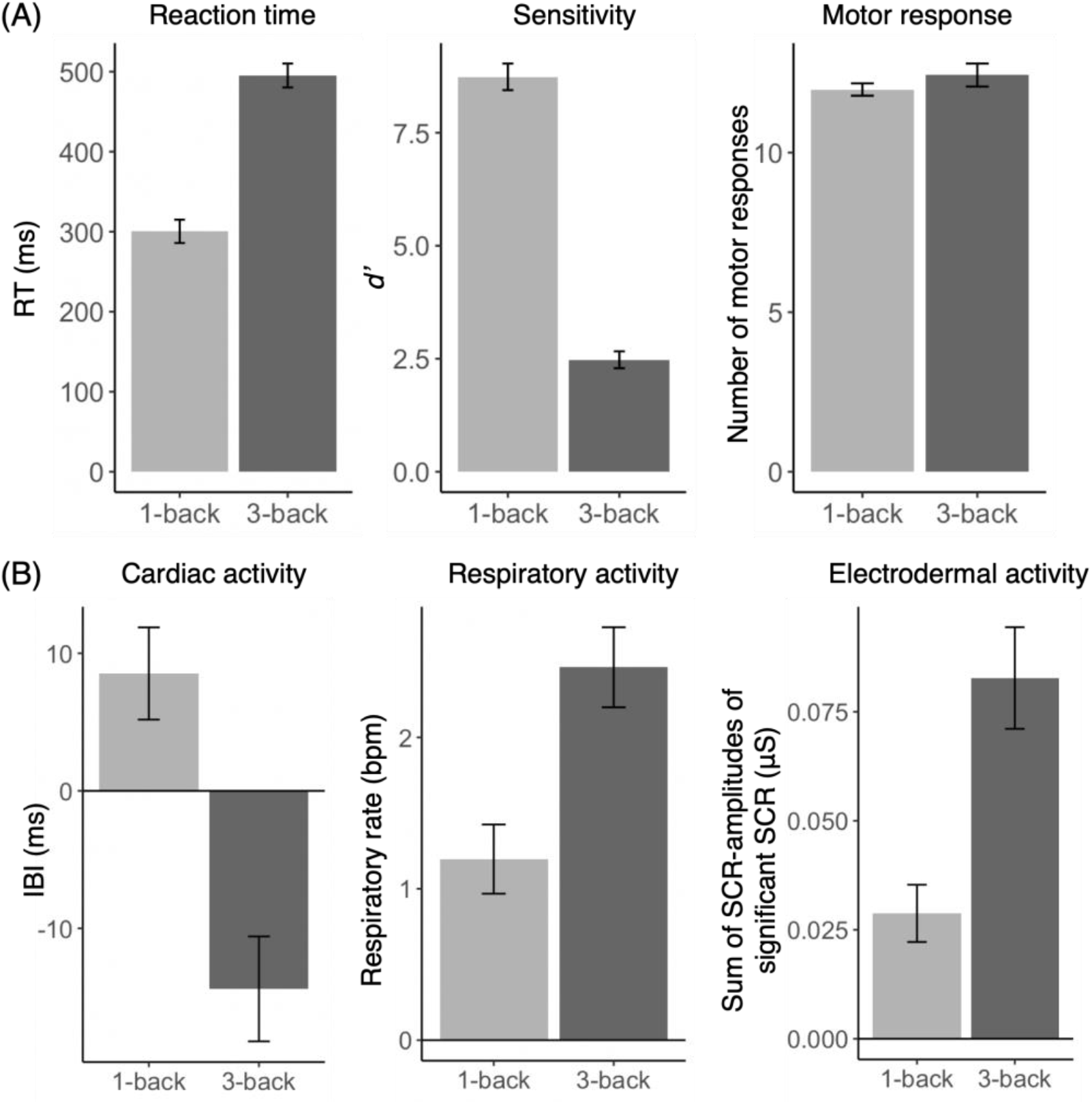
Behavioral and physiological results. (A) Behavioral results. The average reaction time (RT) across participants was 399.10ms (SD = 191.78ms). In 3-back condition, the average RT was 495.19ms (SD = 190.72ms), and in 1-back condition, it was 300.4258ms (SD = 109.87ms), where RT was significantly faster during 1-back compared to 3-back condition (paired t-test: *t*_*(87)*_ = -9.33, *P* < .001). The average sensitivity index (*d’*) across participants was 3.157 (SD = 1.51). In 3-back condition, *d’* was 2.48 (SD = 1.75), and in 1-back condition, it was 8.738 (SD = 2.74), where *d’* was significantly lower during 3-back compared to 1-back condition (*t*_*(87)*_ = 17.97, *P* < .001). The average number of motor responses made overall was 12.21 (SD = 2.72). In 3-back condition, the average number of motor responses made was 12.43 (SD = 3.38), and in 1-back condition, the average number of motor responses made was 11.98 (SD = 1.82). The number of motor responses between the cognitive load conditions does *not* significantly differ (*t*_*(87)*_ = 1.40, *P* > .05). (B) Physiological results. The average change in interbeat interval (IBI) relative to baseline across participants was -2.93ms (SD=29.33ms). In 3-back condition, the average change in IBI was -14.40ms (SD = 28.79ms), and in 1-back condition, it was 8.53ms (SD = 25.30ms). In both conditions, IBI was significantly different from baseline (3-back > baseline: *t*_*(56)*_ = 2.55, *P* < .01; 1-back > baseline: *t*_*(56)*_ = -3.78, *P* < .001). The change in IBI was significantly shorter during 1-back compared to 3-back condition (*t*_*(56)*_ = -5.03, *P* < .001). The average change in respiratory rate relative to baseline across participants was 1.83 breaths per minute (bpm) (SD=1.85bpm). In 3-back condition, the average change in respiratory rate was 2.46bpm (SD = 1.87bpm), and in 1-back condition, it was 1.20bpm (SD = 1.62bpm). In both conditions, respiratory rate was significantly different from baseline (3-back > baseline: *t*_*(49)*_ = -9.32, *P* < .001; 1-back > baseline: *t*_*(49)*_ = 5.23, *P* < .001). The change in respiratory rate was significantly greater during 3-back compared to 1-back condition (*t*_*(49)*_ = 5.16, *P* < .001). The average change in electrodermal activity (EDA), measured by the sum of amplitudes of significant skin conductance response (SCR), relative to baseline across participants was 0.055μS (SD=0.083μS). In 3-back condition, the average change in EDA was 0.083μS (SD = 0.097μS), and in 1-back condition, it was 0.029μS (SD = 0.055μS). In both conditions, EDA was significantly different from baseline (3-back > baseline: *t*_*(68)*_ = 7.10, *P* < .001; 1-back > baseline: *t*_*(68)*_ = 4.38, *P* < .001). The change in EDA was significantly greater during 1-back compared to 3-back condition (*t*_*(68)*_ = 5.25, *P* < .001). Error bars indicate ± 1 standard error across participants. These behavioral and physiological results showed increasing cognitive demand in both conditions, while the 3-back condition showed poorer performance and greater changes in autonomic activity compared to 1-back condition.

**Fig. S2.**
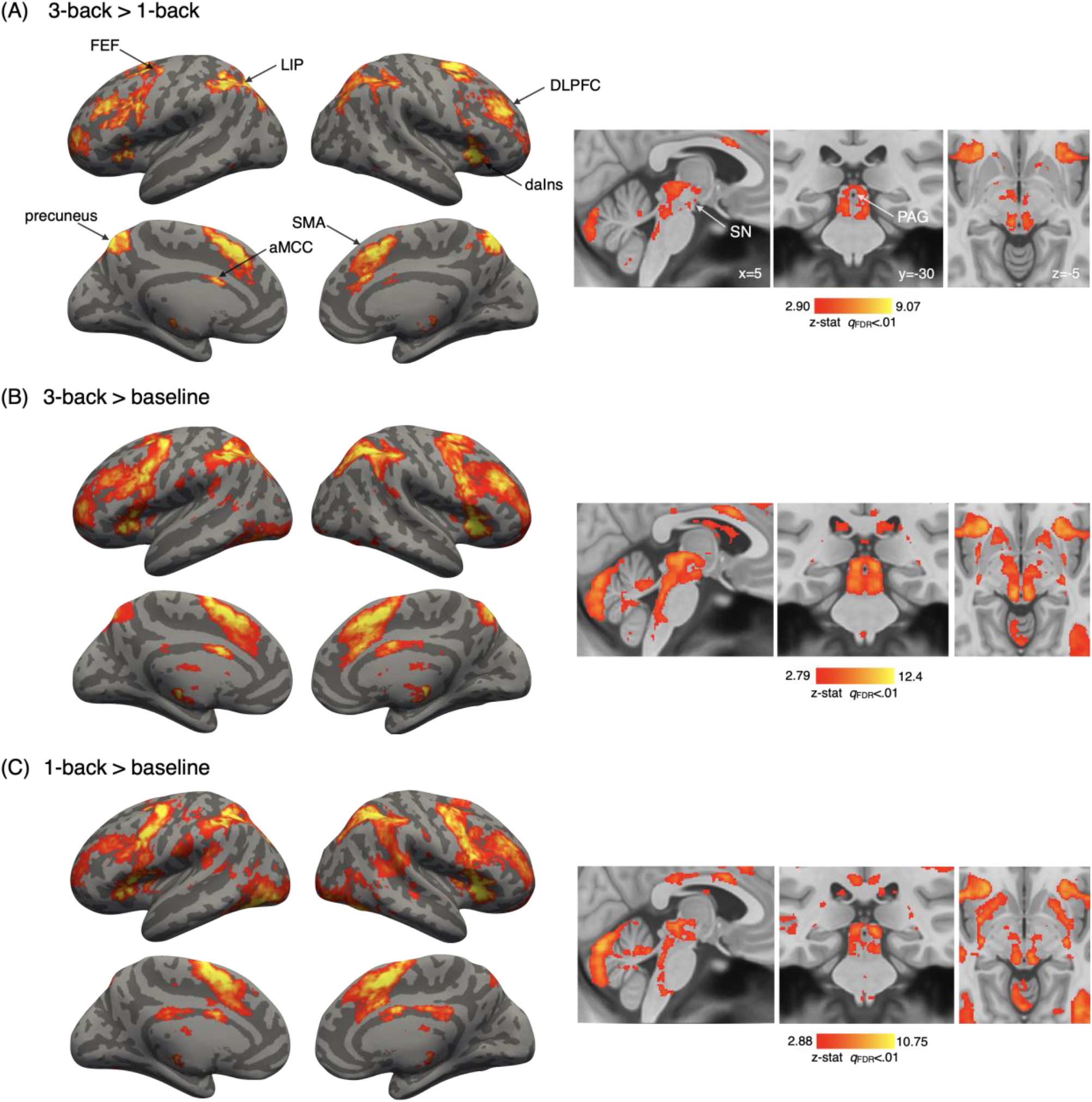
Whole-brain univariate voxel-wise analysis of BOLD fMRI data acquired during an N-back working memory task. Whole-brain contrasts of the 3-back and 1-back conditions identified brain regions that are typically implicated in working memory and visceromotor regulation. These include regions important for executive function, including working memory (dorsolateral prefrontal cortex (DLPFC); dorsal anterior insula (daIns)), top-down control of eye movement (lateral intraparietal cortex (LIP); frontal eye field (FEF)), motor planning (supplementary motor area (SMA); precuneus), and visceromotor control (anterior mid cingulate (aMCC); periaqueductal grey (PAG); substantia nigra (SN); cerebellum) among others (Amiez & Petrides, 2014; Grefkes et al., 2004; Nelson et al., 2010; Owen et al., 2005; Yarkoni et al., 2011).

**Table S1.** Linear mixed effects model summary. F-values and P-values were estimated using Satterthwaite’s method for denominator degrees-of-freedom and F-statistic. For the details of model description, see Online Methods. *** denotes *P* < .001; ** denotes *P* < .01; * denotes *P* < .05.

|  | Sum of squares | Mean square | Numerator df | Denominator df | F value | *P* |
| --- | --- | --- | --- | --- | --- | --- |
| laterality | 32.25 | 32.25 | 1 | 81.680 | 1.7239 | 0.1928628 |
| voxel depth | 277.52 | 277.52 | 1 | 81.782 | 14.8356 | 0.0002321 *** |
| cognitive load | 460.46 | 460.46 | 1 | 77.477 | 24.6150 | 4.05e-06 *** |
| laterality : voxel depth | 26.22 | 26.22 | 1 | 81.318 | 1.4016 | 0.2399106 |
| laterality : cognitive load | 0.56 | 0.56 | 1 | 78.135 | 0.0299 | 0.8631236 |
| voxel depth : cognitive load | 26.83 | 26.83 | 1 | 73.415 | 1.4341 | 0.2349413 |
| laterality : voxel depth : cognitive load | 0.01 | 0.01 | 1 | 77.959 | 0.0003 | 0.9861275 |

**Fig. S3.**
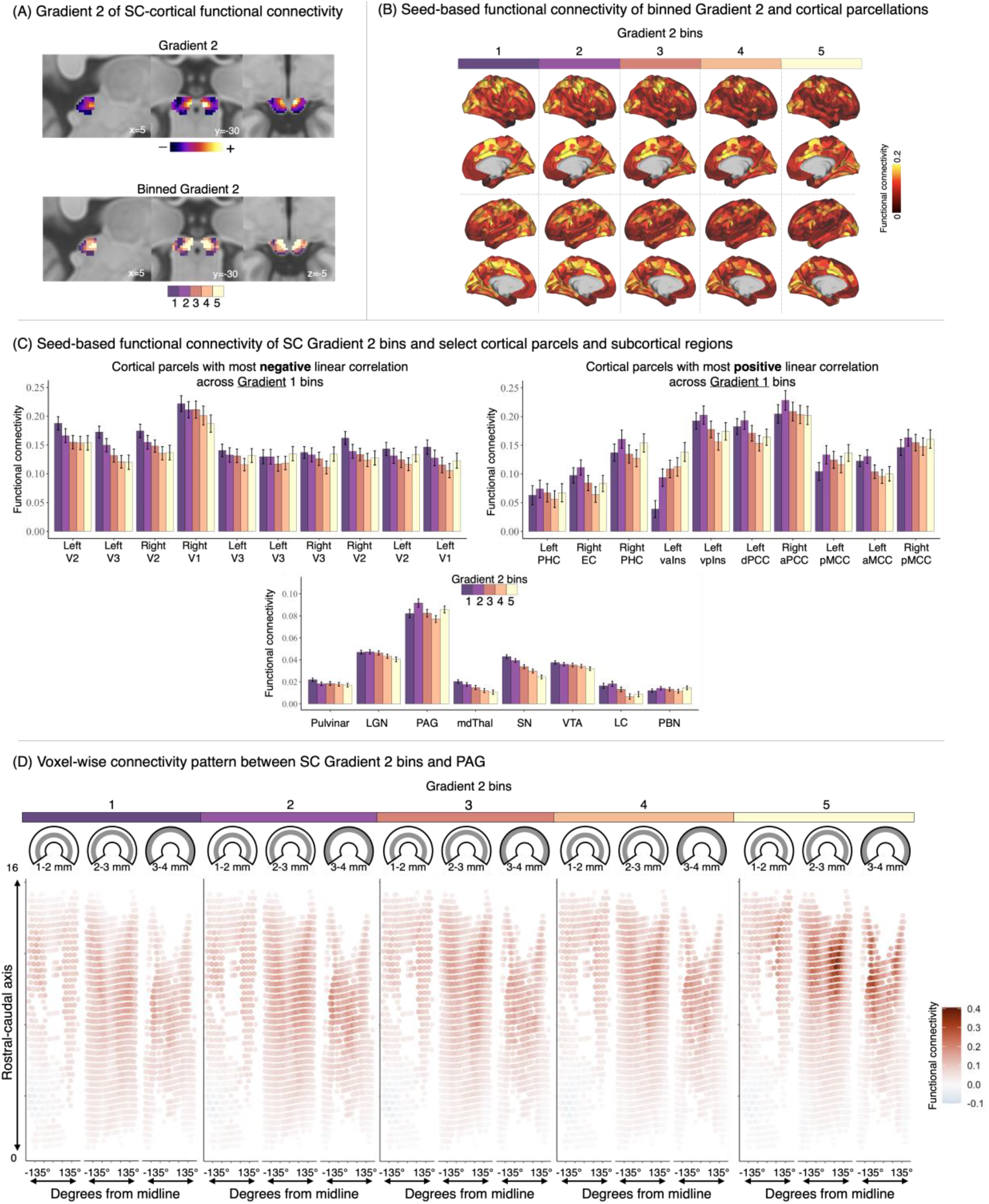
Gradient 2 of SC functional connectivity with the cerebral cortex. (A) Gradient 2 and binned gradient 2 of SC functional connectivity with the cerebral cortex. The transition in functional connectivity pattern was roughly along the medial-lateral axis, but the interpretation remains unclear. (B) Seed-based functional connectivity between SC Gradient 2 bins and cortical parcels. Gradient 2 did not show clear pattern of variation in connectivity pattern across five bins by visual inspection. (C) Seed-based functional connectivity between SC Gradient 2 bins and select cortical parcels and subcortical regions. Cortical parcels were selected based on the rank order of parcels with 10 most negative and 10 most positive linear trends in functional connectivity across Gradient 1 bins. Compared to Gradient 1 results, the same cortical parcels in the visual cortex showed weaker linear decrease in general. Cortical parcels that showed linear increase in Gradient 1, however, did not show clear linear increase in Gradient 2 (except left vaIns). Error bars indicate ± 1 standard error across participants. Compared to Gradient 1 results, the same subcortical regions do not show a clear linear trend with Gradient 2 bins, especially in pulvinar and PAG. Mediodorsal thalamus, substantia nigra, and ventral tegmental area show a negative linear trend across Gradient 2 bins. Error bars indicate ± 1 standard error across participants. (D) Seed-based functional connectivity between SC gradient bins and PAG voxels. Each panel displays three radial cross sections of the PAG cylinder (e.g., 1-2 mm). The y-axis illustrates the rostro-caudal position of the PAG (mm). The x-axis represents +/- 135° from the brain midline. Bin 5 of Gradient 2 showed a similar pattern of strong functional connectivity with dorsolateral PAG as seen in Gradient 1, but more towards the rostral end.

**Fig. S4.**
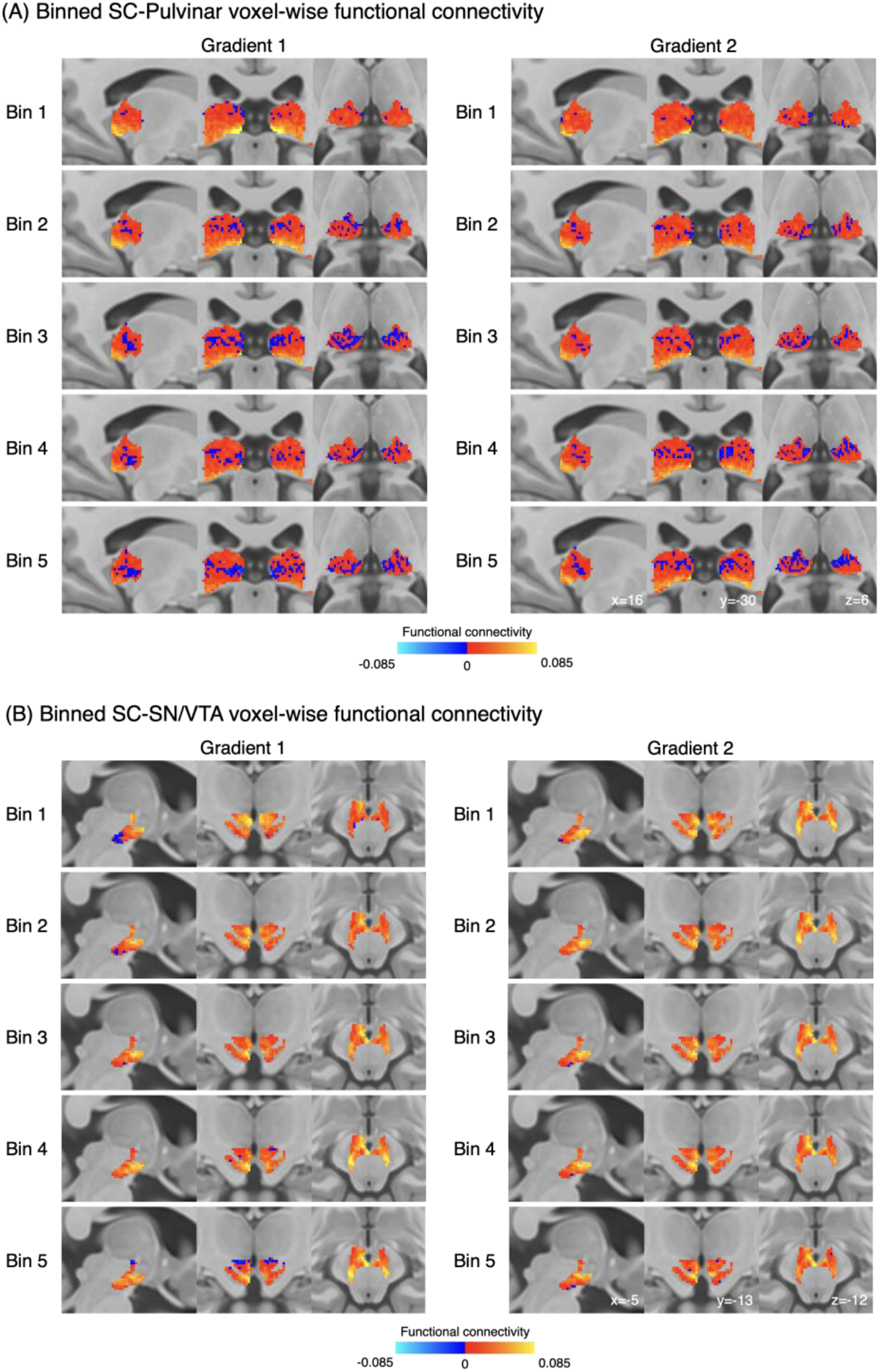
Seed-based functional connectivity between SC gradient bins and each voxel within pulvinar, SN, and VTA. Pulvinar showed widespread positive correlations with bin 1 of Gradient 1 (superficial layers) in its ventroposterior subregion, while this pattern is less strong in other Gradient 1 bins and in the Gradient 2 bins. SN and VTA showed negative correlations with bin 1 of Gradient 1 (near superficial layers) in their ventroposterior subregions and strong positive correlations in their dorsal subregions. This connectivity pattern was altered at bin 5 of Gradient 1 (near deep layers), where the ventroposterior subregions showed positive correlations and the dorsal subregions showed some negative correlations. Connectivity patterns between SN/VTA and Gradient 2 bins did not vary as clearly as in Gradient 1 bins. Abbreviation: SC: superior colliculus; PAG: periaqueductal grey; SN: substantia nigra; VTA: ventral tegmental area.

### Online Methods

#### Participants

One-hundred-and-forty participants were recruited from the Greater Boston area, of which 41 withdrew participation or ended the MRI scan session before starting the working memory task. Eleven participants were excluded due to poor signal quality (e.g., registration-failures caused by participant motion). We additionally screened our participants for excessive head motion (defined as > 0.5 mm framewise displacement in > 20% of TRs per run); none was excluded based on these criteria. The final sample therefore consisted of 88 participants (M_age_ = 27.66 years, SD = 6.18 years, 39 females). All participants were between 18 and 40 years old, were right-handed, had normal or corrected to normal vision, were not pregnant, were fluent English speakers, and had no known neurological or psychiatric illnesses. Participants were excluded from participating in the study if they were claustrophobic or had any metal implants that could cause harm during scanning. All participants provided written informed consent and study procedures were completed as approved by the Partners’ Healthcare Institutional Review Board. Of all included participants, 86 had completed fMRI data for the duration of the task, whereas the other two participants had partial data due to image reconstruction failure for TRs after block 6. To include as much data as possible, we analyzed all usable data from all 88 participants.

#### Experimental paradigm

Participants completed a visual *N*-back working memory task during fMRI scanning, based on the experimental design of prior studies (Gray et al., 2003; van Ast et al., 2016). The task was administered during a single scanning session and pseudorandomly alternated between blocks of either 1-back or 3-back conditions with 10 trials in each block. The session included a total of 12 blocks that were presented in ABBA or BAAB order (counterbalanced across participants) and were each followed by a 25 s rest period. Each block began with a cue indicating the current task (1-back or 3-back). The task was designed with fixed proportions of 20% targets and 80% nontargets (12.5% of which were lures) in each block. Lure trials were defined as 2-back matches in the 1-back blocks and lures in 3-back blocks were either 2 or 4-back matches. The proportion of lure trials was the same for both 1-back and back blocks to equate the requirement for resolving interference. Each block was separated by a 10s rest period.

On every trial, a letter (Q, W, R, S, or T) was presented at the center of the visual field for 2 s followed by a fixation cross for 2 s. Participants were instructed to respond with a button press when the letter on the screen matched the one presented *n* trials ago. The task was administered in MATLAB (RRID:SCR_001622, The MathWorks), using the Psychophysics Toolbox extensions (RRID:SCR_002881; Kleiner, 2007). Stimuli were projected so participants could view them on a mirror fixed to the head coil used for data acquisition. Responses were recorded using an MR-compatible button box. Response times and hit rates for target trials were computed using this signal and served as the primary behavioral outcomes. Before and after the *N*-back task, participants completed a series of self-report items (using a five-point bipolar visual analog scale) to indicate the extent to which they felt awake (vs sleepy), energetic (vs fatigued), engaged (vs bored), pleasant (vs unpleasant), calm (vs tense), and relaxed (vs restless). They were also asked to indicate the extent to which the task would be (was) demanding, whether they would have (had) enough resources to complete the task and whether they would (did) perform well. Before scanning, participants completed a practice session on a laptop. These practice sessions mirrored the task used during scanning, including alternating blocks of one-back and three-back trials, for a total of 48 trials. The goal of the practice session was to ensure that participants were adequately familiar with the N-back task.

#### MRI Data Acquisition and Preprocessing

Gradient-echo echo-planar imaging BOLD-fMRI was performed on a 7-Tesla (7T) Siemens MRI scanner. Functional images were acquired using GRAPPA-EPI sequence: echo time = 28 ms, repetition time = 2.34 s, flip angle = 75°, slice orientation = transversal (axial), anterior to posterior phase encoding, voxel size = 1.1 mm isotropic, gap between slices = 0 mm, number of slices = 123, field of view = 205 × 205 mm^2^, GRAPPA acceleration factor = 3; echo spacing = 0.82 ms, bandwidth = 1414 Hz per pixel, partial Fourier in the phase encode direction = 7/8. A custom-built 32-channel radiofrequency coil head array was used for reception. Radiofrequency transmission was provided by a detunable band-pass birdcage coil.

Structural images were acquired using a T1-weighted EPI sequence, selected so that functional and structural images had similar spatial distortions, which facilitated co-registration and subsequent normalization of data to MNI space. Structural scan parameters were echo time = 22 ms, repetition time = 8.52 s, flip angle = 90°, number of slices = 126, slice orientation = transversal (axial), voxel size = 1.1 mm isotropic, gap between slices = 0 mm, field of view = 205 × 205 mm^2^, GRAPPA acceleration factor = 3; echo spacing = 0.82 ms, bandwidth =1414 Hz per pixel, partial Fourier in the phase encode direction = 6/8.

MRI data were preprocessed using FMRIPREP (Esteban et al., 2019), a Nipype (Gorgolewski et al., 2011) based tool. Spatial normalization of anatomical data to the ICBM 152 Nonlinear Asymmetrical template version 2009c was performed through nonlinear registration with the antsRegistration tool of ANTs v2.1.0 (Avants et al., 2008; Gorgolewski et al., 2011), using brain-extracted versions of both T1w volume and template. Brain tissue segmentation of cerebrospinal fluid (CSF), white-matter (WM) and gray-matter (GM) was performed on the brain-extracted T1w using FAST (FMRIB’s Automated Segmentation Tool; FSL v5.0.9) (Zhang et al., 2001). Functional data were slice time corrected using 3dTshift from AFNI v16.2.07 (Cox, 1996) and motion corrected using MCFLIRT (FSL v5.0.9) (Jenkinson et al., 2002). This was followed by co-registration to the corresponding T1w using boundary-based registration (Greve & Fischl, 2009) with six degrees of freedom, using FLIRT (FMRIB’s Linear Image Registration Tool; FSL v5.0.9) (Jenkinson et al., 2002; Jenkinson & Smith, 2001). Motion correcting transformations, BOLD-to-T1w transformation, and T1w-to-template (MNI) warp were concatenated and applied in a single step using ants ApplyTransforms (ANTs v2.1.0) using Lanczos interpolation. Physiological noise regressors were extracted using the aCompCor method (Muschelli et al., 2014a), taking the top five principle components from subject-specific CSF and WM masks, where the masks were generated by thresholding the WM/CSF masks derived from fast at 99% probability, constraining the CSF mask to the ventricles (using the ALVIN mask; Kempton et al., 2011), and eroding the WM mask using the binary_erosion function in SciPy v.1.6.1 (Virtanen et al., 2020). Frame-wise displacement (Power et al., 2014) was calculated for each functional run using the implementation of Nipype. Many internal operations of fmriprep use Nilearn (Abraham et al., 2014; RRID: SCR_001362), principally within the BOLD-processing workflow. For all participants, the quality of brain masks, tissue segmentation, and MNI registration was visually inspected for errors using the html figures provided by the fmriprep pipeline.

#### SC Localization and Alignment

To account for individual variability in the spatial alignment of the SC to MNI space, we adapted a method from our previous studies of the periaqueductal gray (PAG; Kragel et al., 2019; Satpute et al., 2013) and automated a procedure to anatomically localize the SC in each participant. This procedure began with a hand-drawn SC mask template which was subsequently masked to include voxels in a participant-specific grey and white matter probability masks (> 25%) and exclude voxels in a participant-specific CSF probability mask (> 50%). This mask was further refined to exclude voxels in a participant-specific mask of the PAG and cerebral aqueduct (for procedure, see Kragel et al., 2019), large model residuals from the univariate participant-level analysis (> 85%; see Univariate Analysis below), and any lingering small clusters after the application of other exclusion criteria (< 10 voxels). Participant-specific SC masks were transformed to align all participant whole-brain data in a common space, correcting any misalignment that occurred in the transformation to MNI space. Participant-specific SC masks were aligned using DARTEL (Ashburner, 2007), and the group-level transformations used in this alignment were then applied to participant-specific whole-brain contrast images.

#### SC Layer Estimation

To estimate the location of each voxel in terms of the layer structure of the superior colliculus, we specified two anchor points ventromedial to the SC (right: x=3, y=-30, z=-10; left: x=-3, y=-30, z=-10). These coordinates were manually identified by one of the authors (D.C.). For each side of the superior colliculus, we calculated the Euclidean distance from the anchor to each voxel in the group-level SC mask (for a related approach, see Truong et al., 2020; see also Polimeni et al., 2010). The normalized Euclidean distance was used to approximate the spatial dimension across the SC layers (i.e., voxel depth), such that voxels located in deeper layers had voxel depth closer to 0 with the ventromedial anchors while those in the superficial layers had voxel depth closer to 1.

#### Peripheral physiology recording

All peripheral physiology measures were collected at 1kHz using an AD Instruments Powerlab data acquisition system with MR-compatible sensors and LabChart software (version xx). Data were continuously acquired throughout the entire scan session and partitioned for alignment with fMRI data using experimenter annotations in LabChart and scanner TR events. A piezo-electric pulse transducer (AD Instruments) measured heart rate from the left-hand index fingertip. A respiratory belt with a piezo-electric transducer (UFI) measured respiratory rate and was placed around the lower sternum. Wired Ag/AgCl finger electrodes measured electrodermal activity from sensors containing isotonic paste on the left middle and ring fingertips, with signals amplified by an FE116 GSR amplifier.

Physiological time series data were visually inspected for quality in Biopac Student Lab (version xx) and in custom visualizations using R (v.3.6.2; R Core Team, 2016a) and ggplot2 (v.3.2.1; Wickham, 2009). Time series were classified as clean (i.e. clean of artifacts), noisy (i.e. containing artifacts among periods of usable data), and unusable (i.e. containing artifacts and little to no usable data). Among the 88 participants with usable fMRI data, 57 had clean cardiac data, 50 had clean respiratory data, and 58 had clean electrodermal data. Among electrodermal activity, clean time series contaminated by respiratory noise were included as this noise was irrelevant to the tonic measures of skin conductance used in analysis.

#### Peripheral physiology analysis

Physiological time series data were analyzed using MATLAB toolboxes and custom R scripts. Heart rate was calculated using the PhysIO MATLAB toolbox (Kasper et al., 2017), which used a 0.3 to 9 Hz band-pass filter, and identified cardiac peaks using a two-pass process. The first-pass estimated run average heart rate, detecting peaks exceeding 40% of normalized amplitude, assuming a minimum peak spacing consistent with < 90 beats per minute. First-pass peaks are used to create an averaged template, and the first-pass estimated average heart rate is used as a prior to detect peaks on the second-pass (for more details, see Kasper et al., 2017). PhysIO pipeline peaks were compared with peaks identified in Biopac by trained coders. Discrepancies between the two methods were rare (0.5% of peaks across all runs and participants) and occurred more rarely in the dataset scored by a more experienced coder (0.323% of peaks), compared to the dataset scored by a less experienced coder (0.878% of peaks). Heart rates were smoothed with a 6 second rolling average, downsampled into scan slices, and converted into interbeat intervals. Respiratory rates were calculated using custom R scripts. A 1Hz low-pass filter was applied to respiratory time courses, and local peaks were identified in a sliding 500ms window, removing peaks that fell within 0.5 SD of the run average respiratory value. Respiratory rates were smoothed with a 6 second rolling average and downsampled into scan slices. Tonic electrodermal activity was calculated using Discrete Deconvolution Analysis, as implemented in the Ledalab MATLAB toolbox (Benedek & Kaernbach, 2010).

#### Univariate Analysis

To estimate BOLD activity during the N-back task, a general linear model was applied to preprocessed functional time series (GLM; FSL v5.0.9). First level models were estimated for each participant, including separate regressors for the 3-back and 1-back conditions of each trial (1s). All regressors were convolved with a double gamma hemodynamic function. Regressors were modeled in relation to the implicit baseline of the fixation cross. Nuisance regressors included a run intercept, motion (i.e. translation/rotation in x/y/z planes, and their mean-centered squares, derivatives, and squared derivatives), aCompCor components (5 CSF, 5 WM; Muschelli et al., 2014b), a discrete cosine transformation set with a minimum period of 120 seconds, and spike regressors (> 0.5mm framewise displacement; Satterthwaite et al., 2013). We planned to drop participants from analysis if > 20% of the TRs in their run were excluded via spike regressors, but no participant reached such criterion. Variance inflation factor (VIF) was also estimated to detect multicollinearity, and task regressors with VIF greater than 3 were eliminated from group-level analysis. This resulted in removal of 3-back condition estimates from 11 participants (N = 77) and 1-back estimates from 5 participants (N = 83). No participant had both regressors removed, and in total 72 participants had regressors from both conditions.

First-level whole-brain contrast maps were warped to align SC across participants, then smoothed using a 3 mm FWHM Gaussian kernel and resampled to 0.5 mm voxel size for visualization purposes (Fig. S2 & Fig. 1A). First-level contrast maps were then analyzed at the group level via ordinary least square (OLS) estimation using one-sample *t* tests. Group-level whole-brain contrast results were thresholded by a relatively conservative false discovery rate (*q*_FDR_ < 0.01; compared to the more typical threshold of *q*_FDR_ < 0.05), with a minimum cluster extent of 20 voxels to clean up spurious activation. Group-level voxel-wise SC results were also thresholded by false discovery rate (*q*_FDR_ < 0.01), but no minimum voxel extent was applied.

For the ROI-based analysis of BOLD intensity in the superior colliculus, subject-level superior colliculus-masked contrasts were used as inputs without any smoothing and resampling procedures. We first used paired t-test to compare averaged subject-level SC BOLD intensity during 3-back and 1-back conditions against baseline. We then used a linear mixed effects model to estimate the effects of laterality (left/right), voxel depth (grouped into 10 rank-ordered bins), cognitive load condition (3-back/1-back), their full interactions, and the variability of all fixed effects (i.e., random effects) across participants. By-subject random effects included random intercepts and random slopes for all fixed effects listed above. First-level contrast estimates for each voxel (in total 377 voxels) without smoothing within the group-level SC masks were entered in terms of laterality (2 sides), voxel depth (10 bins), and cognitive loads (2 conditions). In R code (R Core Team, 2016b), estimates were entered into a linear mixed effects model, using the *lmer4* package (Bates et al., 2015), and degrees of freedom were approximated by the Scatterhwaite method, as implemented in the *lmerTest* package (Kuznetsova et al., 2017). The code was entered as follows:

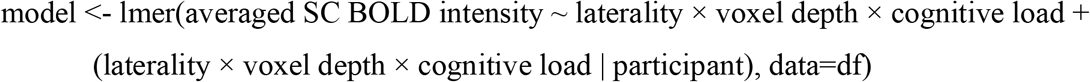

#### Functional Connectivity Gradients Analysis

We derived functional connectivity gradients of the SC using diffusion map embedding (Coifman & Lafon, 2006; Vos de Wael et al., 2020). Diffusion map embedding is a non-linear data dimensionality reduction technique that enables an analysis of similarity structure in functional connectivity patterns by identifying a set of low-dimensional manifolds (i.e., gradients) capturing principal dimensions of spatial variation in connectivity across voxels of interest. As the first step of this analysis, we computed a functional connectivity matrix for each participant between 377 voxels within the group-specific SC mask and 1000 cortical parcels (Schaefer et al., 2018), using the residual BOLD timeseries data generated by the univariate analysis of BOLD data acquired during the N-back task. The residual BOLD timeseries were warped to align SC across participants, the same procedure as that used to generate contrasts maps. This procedure identified four participants whose EPI data had an insufficient coverage of the brain resulting in the number of parcels less than 1000. Therefore, all analyses of functional connectivity gradients were conducted based on the remaining 84 participants. Individual-level functional connectivity matrices were then averaged to yield a group-level matrix.

We applied row-wise thresholding to retain the top 10% connections, with all other connections set to zero. To characterize the similarity structure in functional connectivity between a given pair of a voxel in the SC and a cortical parcel, we converted the thresholded functional connectivity matrix to a non-negative square symmetric affinity matrix expressed as normalized angle similarity. Finally, we used this affinity matrix as input to diffusion map embedding, which yielded 10 gradients identifying the dominant dimensions of spatial variation in the pattern of functional connectivity with the cortex across all voxels in the superior colliculus. In the present study, we focused on the two most dominant gradients in each structure following prior studies of functional connectivity gradients in the brain (Guell et al., 2018; Margulies et al., 2016; Yang et al., 2020). To quantify the relationship between each gradient and the voxel depth of SC layer estimate, we computed voxel-wise Spearman’s rank correlations.

To interpret each gradient of the SC in terms of their connectivity with the rest of the brain, we performed a seed-based functional connectivity analysis. We first discretized each gradient into 5 bins using the percentile ranks of voxel-wise gradient values following previous approaches (Lowe et al., 2019; Royer et al., 2020). This procedure resulted in 5 spatially discontinuous subregions of the SC for each gradient. We calculated the mean timeseries of each gradient bin, which was then used to compute the Pearson’s correlation coefficient with the cortical parcellations for each participant. The group-level connectivity maps were obtained by averaging the correlation data across all participants at each parcellation. To explore the relationship between functional connectivity gradients of the SC and subcortical regions, we performed a similar seed-based analysis of functional connectivity by targeting some structures have been previously identified as closely connected with the superior colliculus, including pulvinar, lateral geniculate nucleus, periaqueductal grey, mediodorsal thalamus, substantia nigra, ventral tegmental area, locus coeruleus, and parabrachial nucleus (for the details of anatomical connectivity evidence, see Benevento & Fallon, 1975; Edwards et al., 1979; Grofová et al., 1978; Redgrave et al., 1987; Sommer & Wurtz, 2004; Waterhouse et al., 1993). These regions have also been found to be functionally important for vision-related processing, visceromotor regulation, and working memory (e.g., Aston-Jones et al., 1996; Davern, 2014; Grieve et al., 2000; Ikemoto et al., 2015; Kragel et al., 2019; Öngür & Price, 2000; Sherman & Koch, 1986; Sotres-Bayón et al., 2001). Anatomical masks for these regions of interest (ROI) were defined based on previous research (see Bianciardi et al., 2016; Cauzzo et al., 2022; Shen et al., 2013; Singh et al., 2020). Averaged functional connectivity was calculated across all voxels within the subcortical regions and then averaged across all functional connectivity values across participants. We additionally investigated mean voxel-wise functional connectivity between each gradient bin and subcortical ROI across all participants.

## Reference

Abraham, A., Pedregosa, F., Eickenberg, M., Gervais, P., Mueller, A., Kossaifi, J., Gramfort, A., Thirion, B., & Varoquaux, G. (2014). Machine learning for neuroimaging with scikit-learn. Frontiers in Neuroinformatics, 8. https://doi.org/10.3389/fninf.2014.00014

Amiez, C., & Petrides, M. (2014). Neuroimaging evidence of the anatomo-functional organization of the human cingulate motor areas. Cerebral Cortex (New York, N.Y.: 1991), 24(3), 563–578. https://doi.org/10.1093/cercor/bhs329

Ashburner, J. (2007). A fast diffeomorphic image registration algorithm. NeuroImage, 38(1), 95–113. https://doi.org/10.1016/j.neuroimage.2007.07.007

Aston-Jones, G., Rajkowski, J., Kubiak, P., Valentino, R. J., & Shipley, M. T. (1996). Chapter 23 Role of the locus coeruleus in emotional activation. In G. Holstege, R. Bandler, & C. B. Saper (Eds.), Progress in Brain Research (Vol. 107, pp. 379–402). Elsevier. https://doi.org/10.1016/S0079-6123(08)61877-4

Avants, B. B., Epstein, C. L., Grossman, M., & Gee, J. C. (2008). Symmetric diffeomorphic image registration with cross-correlation: Evaluating automated labeling of elderly and neurodegenerative brain. Medical Image Analysis, 12(1), 26–41. https://doi.org/10.1016/j.media.2007.06.004

Basso, M. A., Bickford, M. E., & Cang, J. (2021). Unraveling circuits of visual perception and cognition through the superior colliculus. Neuron, 109(6), 918–937. https://doi.org/10.1016/j.neuron.2021.01.013

Bates, D., Mächler, M., Bolker, B., & Walker, S. (2015). Fitting Linear Mixed-Effects Models Using lme4. Journal of Statistical Software, 67(1), Article 1. https://doi.org/10.18637/jss.v067.i01

Benedek, M., & Kaernbach, C. (2010). Decomposition of skin conductance data by means of nonnegative deconvolution. Psychophysiology, 47(4), 647–658. https://doi.org/10.1111/j.1469-8986.2009.00972.x

Benevento, L. A., & Fallon, J. H. (1975). The ascending projections of the superior colliculus in the rhesus monkey (Macaca mulatta). Journal of Comparative Neurology, 160(3), 339–361. https://doi.org/10.1002/cne.901600306

Bianciardi, M., Toschi, N., Eichner, C., Polimeni, J. R., Setsompop, K., Brown, E. N., Hämäläinen, M. S., Rosen, B. R., & Wald, L. L. (2016). In vivo functional connectome of human brainstem nuclei of the ascending arousal, autonomic, and motor systems by high spatial resolution 7-Tesla fMRI. Magma (New York, N.Y.), 29(3), 451–462. https://doi.org/10.1007/s10334-016-0546-3

Bittencourt, A. S., Carobrez, A. P., Zamprogno, L. P., Tufik, S., & Schenberg, L. C. (2004). Organization of single components of defensive behaviors within distinct columns of periaqueductal gray matter of the rat: Role of N-METHYL-d-aspartic acid glutamate receptors. Neuroscience, 125(1), 71–89. https://doi.org/10.1016/j.neuroscience.2004.01.026

Carrive, P., & Morgan, M. M. (2012). Periaqueductal Gray. In The Human Nervous System (pp. 367–400). Elsevier. https://doi.org/10.1016/B978-0-12-374236-0.10010-0

Cauzzo, S., Singh, K., Stauder, M., García-Gomar, M. G., Vanello, N., Passino, C., Staab, J., Indovina, I., & Bianciardi, M. (2022). Functional connectome of brainstem nuclei involved in autonomic, limbic, pain and sensory processing in living humans from 7 Tesla resting state fMRI. NeuroImage, 250, 118925. https://doi.org/10.1016/j.neuroimage.2022.118925

Coifman, R. R., & Lafon, S. (2006). Diffusion maps. Applied and Computational Harmonic Analysis, 21(1), 5–30. https://doi.org/10.1016/j.acha.2006.04.006

Cox, R. W. (1996). AFNI: Software for Analysis and Visualization of Functional Magnetic Resonance Neuroimages. Computers and Biomedical Research, 29(3), 162–173. https://doi.org/10.1006/cbmr.1996.0014

Davern, P. J. (2014). A role for the lateral parabrachial nucleus in cardiovascular function and fluid homeostasis. Frontiers in Physiology, 5. https://www.frontiersin.org/articles/10.3389/fphys.2014.00436

Edwards, S. B., Ginsburgh, C. L., Henkel, C. K., & Stein, B. E. (1979). Sources of subcortical projections to the superior colliculus in the cat. The Journal of Comparative Neurology, 184(2), 309–329. https://doi.org/10.1002/cne.901840207

Esteban, O., Markiewicz, C. J., Blair, R. W., Moodie, C. A., Isik, A. I., Erramuzpe, A., Kent, J. D., Goncalves, M., DuPre, E., Snyder, M., Oya, H., Ghosh, S. S., Wright, J., Durnez, J., Poldrack, R. A., & Gorgolewski, K. J. (2019). fMRIPrep: A robust preprocessing pipeline for functional MRI. Nature Methods, 16(1), 111–116. https://doi.org/10.1038/s41592-018-0235-4

Gandhi, N. J., & Katnani, H. A. (2011). Motor Functions of the Superior Colliculus. Annual Review of Neuroscience, 34, 205–231. https://doi.org/10.1146/annurev-neuro-061010-113728

Gorgolewski, K., Burns, C. D., Madison, C., Clark, D., Halchenko, Y. O., Waskom, M. L., & Ghosh, S. S. (2011). Nipype: A Flexible, Lightweight and Extensible Neuroimaging Data Processing Framework in Python. Frontiers in Neuroinformatics, 0. https://doi.org/10.3389/fninf.2011.00013

Gray, J. R., Chabris, C. F., & Braver, T. S. (2003). Neural mechanisms of general fluid intelligence. Nature Neuroscience, 6(3), 316–322. https://doi.org/10.1038/nn1014

Grefkes, C., Ritzl, A., Zilles, K., & Fink, G. R. (2004). Human medial intraparietal cortex subserves visuomotor coordinate transformation. NeuroImage, 23(4), 1494–1506. https://doi.org/10.1016/j.neuroimage.2004.08.031

Greve, D. N., & Fischl, B. (2009). Accurate and robust brain image alignment using boundary-based registration. NeuroImage, 48(1), 63–72. https://doi.org/10.1016/j.neuroimage.2009.06.060

Grieve, K. L., Acuña, C., & Cudeiro, J. (2000). The primate pulvinar nuclei: Vision and action. Trends in Neurosciences, 23(1), 35–39. https://doi.org/10.1016/S0166-2236(99)01482-4

Grofová, I., Ottersen, O. P., & Rinvik, E. (1978). Mesencephalic and diencephalic afferents to he superior colliculus and periaqueductal gray substance demonstrated by retrograde axonal transport of horseradish peroxidase in the cat. Brain Research, 146(2), 205–220. https://doi.org/10.1016/0006-8993(78)90969-1

Guell, X., Schmahmann, J. D., Gabrieli, J. D., & Ghosh, S. S. (2018). Functional gradients of the cerebellum. ELife, 7, e36652. https://doi.org/10.7554/eLife.36652

Hong, S.-J., Xu, T., Nikolaidis, A., Smallwood, J., Margulies, D. S., Bernhardt, B., Vogelstein, J., & Milham, M. P. (2020). Toward a connectivity gradient-based framework for reproducible biomarker discovery. NeuroImage, 223, 117322. https://doi.org/10.1016/j.neuroimage.2020.117322

Ikemoto, S., Yang, C., & Tan, A. (2015). Basal ganglia circuit loops, dopamine and motivation: A review and enquiry. Behavioural Brain Research, 290, 17–31. https://doi.org/10.1016/j.bbr.2015.04.018

Isa, T., Marquez-Legorreta, E., Grillner, S., & Scott, E. K. (2021). The tectum/superior colliculus as the vertebrate solution for spatial sensory integration and action. Current Biology, 31(11), R741–R762. https://doi.org/10.1016/j.cub.2021.04.001

Jenkinson, M., Bannister, P., Brady, M., & Smith, S. (2002). Improved Optimization for the Robust and Accurate Linear Registration and Motion Correction of Brain Images. NeuroImage, 17(2), 825–841. https://doi.org/10.1006/nimg.2002.1132

Jenkinson, M., & Smith, S. (2001). A global optimisation method for robust affine registration of brain images. Medical Image Analysis, 5(2), 143–156. https://doi.org/10.1016/S1361-8415(01)00036-6

Jun, E. J., Bautista, A. R., Nunez, M. D., Allen, D. C., Tak, J. H., Alvarez, E., & Basso, M. A. (2021). Causal role for the primate superior colliculus in the computation of evidence for perceptual decisions. Nature Neuroscience, 24(8), 1121–1131. https://doi.org/10.1038/s41593-021-00878-6

Kasper, L., Bollmann, S., Diaconescu, A. O., Hutton, C., Heinzle, J., Iglesias, S., Hauser, T. U., Sebold, M., Manjaly, Z.-M., Pruessmann, K. P., & Stephan, K. E. (2017). The PhysIO Toolbox for Modeling Physiological Noise in fMRI Data. Journal of Neuroscience Methods, 276, 56–72. https://doi.org/10.1016/j.jneumeth.2016.10.019

Katsumi, Y., Theriault, J. E., Quigley, K., & Barrett, L. F. (2021). Allostasis as a core feature of hierarchical gradients in the human brain [Preprint]. PsyArXiv. https://doi.org/10.31234/osf.io/wezv8

Kempton, M. J., Underwood, T. S. A., Brunton, S., Stylios, F., Schmechtig, A., Ettinger, U., Smith, M. S., Lovestone, S., Crum, W. R., Frangou, S., Williams, S. C. R., & Simmons, A. (2011). A comprehensive testing protocol for MRI neuroanatomical segmentation techniques: Evaluation of a novel lateral ventricle segmentation method. NeuroImage, 58(4), 1051–1059. https://doi.org/10.1016/j.neuroimage.2011.06.080

Kleiner, M. (2007). What’s new in Psychtoolbox-3? 89.

Kragel, P. A., Bianciardi, M., Hartley, L., Matthewson, G., Choi, J.-K., Quigley, K. S., Wald, L. L., Wager, T. D., Barrett, L. F., & Satpute, A. B. (2019). Functional Involvement of Human Periaqueductal Gray and Other Midbrain Nuclei in Cognitive Control. Journal of Neuroscience, 39(31), 6180–6189. https://doi.org/10.1523/JNEUROSCI.2043-18.2019

Krebs, R. M., Woldorff, M. G., Tempelmann, C., Bodammer, N., Noesselt, T., Boehler, C. N., Scheich, H., Hopf, J.-M., Duzel, E., Heinze, H.-J., & Schoenfeld, M. A. (2010). High-Field fMRI Reveals Brain Activation Patterns Underlying Saccade Execution in the Human Superior Colliculus. PLOS ONE, 5(1), e8691. https://doi.org/10.1371/journal.pone.0008691

Kuznetsova, A., Brockhoff, P. B., & Christensen, R. H. B. (2017). lmerTest Package: Tests in Linear Mixed Effects Models. Journal of Statistical Software, 82(1), Article 1. https://doi.org/10.18637/jss.v082.i13

Liu, X., Huang, H., Snutch, T. P., Cao, P., Wang, L., & Wang, F. (2022). The Superior Colliculus: Cell Types, Connectivity, and Behavior. Neuroscience Bulletin. https://doi.org/10.1007/s12264-022-00858-1

Loureiro, J. R., Hagberg, G. E., Ethofer, T., Erb, M., Bause, J., Ehses, P., Scheffler, K., & Himmelbach, M. (2017). Depth-dependence of visual signals in the human superior colliculus at 9.4 T. Human Brain Mapping, 38(1), 574–587. https://doi.org/10.1002/hbm.23404

Lovejoy, L. P., & Krauzlis, R. J. (2010). Inactivation of primate superior colliculus impairs covert selection of signals for perceptual judgments. Nature Neuroscience, 13(2), Article 2. https://doi.org/10.1038/nn.2470

Lowe, A. J., Paquola, C., Wael, R. V. de, Girn, M., Lariviere, S., Tavakol, S., Caldairou, B., Royer, J., Schrader, D. V., Bernasconi, A., Bernasconi, N., Spreng, R. N., & Bernhardt, A.C. (2019). Targeting age-related differences in brain and cognition with multimodal imaging and connectome topography profiling. Human Brain Mapping, 40(18), 5213–5230. https://doi.org/10.1002/hbm.24767

Mandrick, K., Peysakhovich, V., Rémy, F., Lepron, E., & Causse, M. (2016). Neural and psychophysiological correlates of human performance under stress and high mental workload. Biological Psychology, 121, 62–73. https://doi.org/10.1016/j.biopsycho.2016.10.002

Margulies, D. S., Ghosh, S. S., Goulas, A., Falkiewicz, M., Huntenburg, J. M., Langs, G., Bezgin, G., Eickhoff, S. B., Castellanos, F. X., Petrides, M., Jefferies, E., & Smallwood, J. (2016). Situating the default-mode network along a principal gradient of macroscale cortical organization. Proceedings of the National Academy of Sciences, 113(44), 12574–12579. https://doi.org/10.1073/pnas.1608282113

May, P. J. (2006). The mammalian superior colliculus: Laminar structure and connections. In J. A. Büttner-Ennever (Ed.), Progress in Brain Research (Vol. 151, pp. 321–378). Elsevier. https://doi.org/10.1016/S0079-6123(05)51011-2

Muschelli, J., Nebel, M. B., Caffo, B. S., Barber, A. D., Pekar, J. J., & Mostofsky, S. H. (2014a). Reduction of motion-related artifacts in resting state fMRI using aCompCor. NeuroImage, 96, 22–35. https://doi.org/10.1016/j.neuroimage.2014.03.028

Muschelli, J., Nebel, M. B., Caffo, B. S., Barber, A. D., Pekar, J. J., & Mostofsky, S. H. (2014b). Reduction of motion-related artifacts in resting state fMRI using aCompCor. NeuroImage, 96, 22–35. https://doi.org/10.1016/j.neuroimage.2014.03.028

Nelson, S. M., Dosenbach, N. U. F., Cohen, A. L., Wheeler, M. E., Schlaggar, B. L., & Petersen, S. E. (2010). Role of the anterior insula in task-level control and focal attention. Brain Structure and Function, 214(5), 669–680. https://doi.org/10.1007/s00429-010-0260-2

Öngür, D., & Price, J. L. (2000). The Organization of Networks within the Orbital and Medial Prefrontal Cortex of Rats, Monkeys and Humans. Cerebral Cortex, 10(3), 206–219. https://doi.org/10.1093/cercor/10.3.206

Owen, A. M., McMillan, K. M., Laird, A. R., & Bullmore, E. (2005). N-back working memory paradigm: A meta-analysis of normative functional neuroimaging studies. Human Brain Mapping, 25(1), 46–59. https://doi.org/10.1002/hbm.20131

Polimeni, J. R., Greve, D. N., Fischl, B., & Wald, L. L. (2010). Depth-resolved laminar analysis of resting-state fluctuation amplitude in high-resolution 7T fMRI. 18th Annual Meeting International Society for Magnetic Resonance in Medicine, Stockholm, 1168.

Power, J. D., Mitra, A., Laumann, T. O., Snyder, A. Z., Schlaggar, B. L., & Petersen, S. E. (2014). Methods to detect, characterize, and remove motion artifact in resting state fMRI. NeuroImage, 84, 320–341. https://doi.org/10.1016/j.neuroimage.2013.08.048

R Core Team. (2016a). R: A language and environment for statistical computing. R Foundation for Statistical Computing. https://www.R-project.org

R Core Team. (2016b). R: A language and environment for statistical computing. 16.

Redgrave, P., Mitchell, I. J., & Dean, P. (1987). Descending projections from the superior colliculus in rat: A study using orthograde transport of wheatgerm-agglutinin conjugated horseradish peroxidase. Experimental Brain Research, 68(1), 147–167. https://doi.org/10.1007/BF00255241

Royer, J., Paquola, C., Larivière, S., Vos de Wael, R., Tavakol, S., Lowe, A. J., Benkarim, O., Evans, A. C., Bzdok, D., Smallwood, J., Frauscher, B., & Bernhardt, B. C. (2020). Myeloarchitecture gradients in the human insula: Histological underpinnings and association to intrinsic functional connectivity. NeuroImage, 216, 116859. https://doi.org/10.1016/j.neuroimage.2020.116859

Satpute, A. B., Wager, T. D., Cohen-Adad, J., Bianciardi, M., Choi, J.-K., Buhle, J. T., Wald, L. L., & Barrett, L. F. (2013). Identification of discrete functional subregions of the human periaqueductal gray. Proceedings of the National Academy of Sciences, 110(42), 17101–17106. https://doi.org/10.1073/pnas.1306095110

Satterthwaite, T. D., Elliott, M. A., Gerraty, R. T., Ruparel, K., Loughead, J., Calkins, M. E., Eickhoff, S. B., Hakonarson, H., Gur, R. C., Gur, R. E., & Wolf, D. H. (2013). An improved framework for confound regression and filtering for control of motion artifact in the preprocessing of resting-state functional connectivity data. NeuroImage, 64, 240–256. https://doi.org/10.1016/j.neuroimage.2012.08.052

Schaefer, A., Kong, R., Gordon, E. M., Laumann, T. O., Zuo, X.-N., Holmes, A. J., Eickhoff, S. B., & Yeo, B. T. T. (2018). Local-Global Parcellation of the Human Cerebral Cortex from Intrinsic Functional Connectivity MRI. Cerebral Cortex, 28(9), 3095–3114. https://doi.org/10.1093/cercor/bhx179

Shen, X., Tokoglu, F., Papademetris, X., & Constable, R. T. (2013). Groupwise whole-brain parcellation from resting-state fMRI data for network node identification. NeuroImage, 82, 403–415. https://doi.org/10.1016/j.neuroimage.2013.05.081

Sherman, S. M., & Koch, C. (1986). The control of retinogeniculate transmission in the mammalian lateral geniculate nucleus. Experimental Brain Research, 63(1), 1–20. https://doi.org/10.1007/BF00235642

Singh, K., Indovina, I., Augustinack, J. C., Nestor, K., García-Gomar, M. G., Staab, J. P., & Bianciardi, M. (2020). Probabilistic Template of the Lateral Parabrachial Nucleus, Medial Parabrachial Nucleus, Vestibular Nuclei Complex, and Medullary Viscero-Sensory-Motor Nuclei Complex in Living Humans From 7 Tesla MRI. Frontiers in Neuroscience, 13. https://www.frontiersin.org/article/10.3389/fnins.2019.01425

Sommer, M. A., & Wurtz, R. H. (2004). What the Brain Stem Tells the Frontal Cortex. I. Oculomotor Signals Sent From Superior Colliculus to Frontal Eye Field Via Mediodorsal Thalamus. Journal of Neurophysiology, 91(3), 1381–1402. https://doi.org/10.1152/jn.00738.2003

Sotres-Bayón, F., Torres-López, E., López-Ávila, A., del Ángel, R., & Pellicer, F. (2001). Lesion and electrical stimulation of the ventral tegmental area modify persistent nociceptive behavior in the rat. Brain Research, 898(2), 342–349. https://doi.org/10.1016/S0006-8993(01)02213-2

Truong, P., Kim, J. H., Savjani, R., Sitek, K. R., Hagberg, G. E., Scheffler, K., & Ress, D. (2020). Depth relationships and measures of tissue thickness in dorsal midbrain. Human Brain Mapping, 41(18), 5083–5096. https://doi.org/10.1002/hbm.25185

van Ast, V. A., Spicer, J., Smith, E. E., Schmer-Galunder, S., Liberzon, I., Abelson, J. L., & Wager, T. D. (2016). Brain Mechanisms of Social Threat Effects on Working Memory. Cerebral Cortex (New York, N.Y.: 1991), 26(2), 544–556. https://doi.org/10.1093/cercor/bhu206

Virtanen, P., Gommers, R., Oliphant, T. E., Haberland, M., Reddy, T., Cournapeau, D., Burovski, E., Peterson, P., Weckesser, W., Bright, J., van der Walt, S. J., Brett, M., Wilson, J., Millman, K. J., Mayorov, N., Nelson, A. R. J., Jones, E., Kern, R., Larson, E., … van Mulbregt, P. (2020). SciPy 1.0: Fundamental algorithms for scientific computing in Python. Nature Methods, 17(3), 261–272. https://doi.org/10.1038/s41592-019-0686-2

Vos de Wael, R., Benkarim, O., Paquola, C., Lariviere, S., Royer, J., Tavakol, S., Xu, T., Hong, S.-J., Langs, G., Valk, S., Misic, B., Milham, M., Margulies, D., Smallwood, J., & Bernhardt, B. C. (2020). BrainSpace: A toolbox for the analysis of macroscale gradients in neuroimaging and connectomics datasets. Communications Biology, 3(1), Article 1. https://doi.org/10.1038/s42003-020-0794-7

Wang, Y. C., Bianciardi, M., Chanes, L., & Satpute, A. B. (2020). Ultra High Field fMRI of Human Superior Colliculi Activity during Affective Visual Processing. Scientific Reports, 10(1), 1331. https://doi.org/10.1038/s41598-020-57653-z

Waterhouse, B. D., Border, B., Wahl, L., & Mihailoff, G. A. (1993). Topographic organization of rat locus coeruleus and dorsal raphe nuclei: Distribution of cells projecting to visual system structures. Journal of Comparative Neurology, 336(3), 345–361. https://doi.org/10.1002/cne.903360304

Wickham, H. (2009). ggplot2: Elegant Graphics for Data Analysis. Springer-Verlag. https://doi.org/10.1007/978-0-387-98141-3

Yang, S., Meng, Y., Li, J., Li, B., Fan, Y.-S., Chen, H., & Liao, W. (2020). The thalamic functional gradient and its relationship to structural basis and cognitive relevance. NeuroImage, 218, 116960. https://doi.org/10.1016/j.neuroimage.2020.116960

Yarkoni, T., Poldrack, R. A., Nichols, T. E., Van Essen, D. C., & Wager, T. D. (2011). Large-scale automated synthesis of human functional neuroimaging data. Nature Methods, 8(8), 665–670. https://doi.org/10.1038/nmeth.1635

Zhang, Y., Brady, M., & Smith, S. (2001). Segmentation of brain MR images through a hidden Markov random field model and the expectation-maximization algorithm. IEEE Transactions on Medical Imaging, 20(1), 45–57. https://doi.org/10.1109/42.906424

